# Ancestral persistence of vestibulo-spinal reflexes in axial muscles in humans

**DOI:** 10.1101/564617

**Authors:** Etienne Guillaud, Céline Faure, Emilie Doat, Laurent Bouyer, Dominique Guehl, Jean-René Cazalets

## Abstract

Accurate control of the trunk is essential for maintaining balance in upright subjects. Most studies addressing vestibulo-spinal reflexes have investigated the role of the lower limbs, while limited attention has been paid to the back muscles. To address this issue, we challenged the persistence of vestibular evoked myogenic potentials (VEMPs) in back muscles in situations in which the leg muscle responses were modulated. Nineteen subjects were submitted to galvanic vestibular stimulations (GVS). Body sway and VEMPs were recorded in the paraspinal and limb muscles. During treadmill locomotion, the VEMPS in the lower limbs were observed only during the stance phase, whereas the axial VEMPs were observed during all phases. In upright standing subjects, slight head contact was sufficient to abolish the VEMPs in the lower limbs, while the VEMPs remained present in the paraspinal muscles. Similarly, during parabolic flight-induced microgravity, the VEMPs in the lower limb muscles were suppressed, while those in the axial muscles persisted despite the absence of gravitational information from the otolithic system. Our results depict a differentiated control mechanism of axial and appendicular muscles when a perturbation is detected by vestibular inputs. The persistent feature of axial myogenic adjustments suggests that a hard-wired reflex is functionally efficient to maintain posture. By contrast, the ankle responses to perturbations occur only when the accompanying sensory feedback is congruent, challenging the balance task and gravity. Overall, this study using GVS in microgravity is the first to present an approach delineating feed-forward vestibular control in the absence of all feedback.

**NEW AND NOTEWORTHY:** We addressed the degree to which the underlying neural circuitry and mechanisms involved in trunk muscle control have been preserved in human. We highlighted the persistence of vestibular-evoked responses in trunk muscles using galvanic vestibular stimulation, in conditions where leg muscle responses were modulated (walking, standing, microgravity). This suggest a phylogenetically conserved blueprint of sensorimotor organization, with strongly hard-wired vestibulo-spinal inputs to axial motoneurons and a higher degree of flexibility for the late emerging limb.

## INTRODUCTION

During evolution, locomotion has adapted from propulsion by undulation to quadrupedal and finally bipedal propulsion. Despite its decisive advantages, an erect posture strengthens the difficulty of maintaining balance. This ability to adjust the body orientation, particularly during dynamic tasks, highly relies on the vestibular system. This sensorial input provides adequate information related to head movement and orientation and may evoke responses in limb muscles and postural adjustments (Britton et al., 1993; Fitzpatrick et al., 1994). Despite their high degree of automaticity, vestibulomotor responses are extremely flexible and may vary in a task-dependent manner. Various studies have demonstrated that motor responses in the upper and lower limbs following the vestibular detection of motion directly depend on limb engagement in the current task. For instance, using galvanic vestibular stimulation (GVS, Goldberg et al., 1984; Fitzpatrick and Day 2004), arm motor responses were observed during reaching (Bresciani et al., 2002; Mars et al., 2003; Smith and Reynolds, 2017) or body stabilization when the hand is used to help maintain balance (Britton et al., 1993). This response was absent in the contralateral arm, which was not engaged in the task (Britton et al. 1993). Comparable results have been observed in the lower limb muscles since in seated subjects or subjects standing upright while bearing a support, vestibular evoked responses were no longer present in the leg muscles (Fitzpatrick et al., 1994). A recent study (Forbes et al., 2016) showed that a reverse relationship exists between balancing motor command and associated vestibular sensory feedback in a task conducted in the reverse direction of the vestibular evoked compensatory response. The flexibility of vestibulo-spinal processes has also been documented during walking (Bent et al., 2004; Iles et al., 2007; Forbes et al., 2017; Blouin et al. 2011). Altogether, these findings suggest that new relationships between vestibular signals and consequent motor commands can be established according to the context, even in similar tasks.

However, most studies addressing the control of balance in upright subjects have mainly investigated the role of the lower limbs, while limited attention has been paid to the vestibulo-spinal reflexes that occur in the back muscles. This is surprising because the trunk, which represents more than 60% of the total body mass and is located above the center of gravity, is a potential source of major instability. Because this body part is highly articulated and actuated by many muscles, it should be accurately controlled during displacement. Ali and colleagues (2003) recorded functional vestibular evoked responses in the erector spinae muscles in both sitting and upright postures, and any response was measured in the limb muscles in the sitting posture. Interestingly, the subjects’ responses in the erector spinae recorded at L3/L4 displayed latencies of 59 ms, while the latencies in the lower limb muscles were much longer (≈85 ms). These results suggest that muscles of the axial trunk and leg may be controlled by different vestibulo-spinal mechanisms.

Therefore, in this paper, our objective was to challenge the persistence of vestibular evoked responses in the back muscles under conditions in which the lower-body muscle responses are modulated by a task and/or context using GVS. In the first experiment, we compared the GVS phase effect dependency on back muscles and lower limb muscles during walking. In the second experiment, we tested two conditions in which vestibular evoked myogenic potentials (VEMPs) were suppressed in the lower limbs, i.e., in upright subjects bearing a support and upright subjects sustained in a harness. In the third experiment, which represented our “most challenging” condition, we tested the persistence of vestibulo-spinal reflexes in floating subjects in the absence of gravity (during parabolic flight) as no postural adjustments were required. To the best of our knowledge, this study represents the first time GVS responses were studied in microgravity. Altogether, we demonstrate that under all these contextual and/or gravitational conditions, balance and posture were persistently controlled using back muscles whereas the lower limb responses could be flexibly modulated.

## MATERIAL AND METHODS

Nineteen adults participated in this study. The subjects provided written informed consent prior to participating in each experiment. The procedures conformed to the Declaration of Helsinki and were approved by the French National Research Ethics Committee (Comité de Protection des Personnes Sud-Ouest et Outre Mer III) under agreement number 2011-A00424-37. In all experiments, we used similar tools and methods, including pulse GVS stimulation, electromyography and 3D kinematics.

### Vestibular stimulations

Bipolar binaural galvanic electrical stimulations were applied to the mastoid processes. The electrodes were circular with a contact surface of 9 cm^2^ (Axelgaard Pals Platinium, Ø32 mm). Square pulse stimulations (3.5 mA; 175 ms) were delivered using an isolated constant current stimulator (Digitimer DS-5, CE mark certification for medical devices) with randomized and unpredictable delays (5 s minimum between two stimulations).

### Electromyographic recordings and analysis

EMGs were recorded bilaterally on the *medial gastrocnemius*, *anterior tibialis*, and *erector spinae* at the C7, T7 and L4 levels (De Sèze et al., 2008; Cecatto et al., 2009). During walking on a treadmill, we used a wireless EMG system (Kine ehf, Iceland; 1.562 KHz, x10,000 amplifier), while all other recordings were performed using an analogical amplifier (TeleEMG, BTS, Milano, Italy; x1000 amplifier) linked to an ITC-18 A/D interface (Heka, Lambrecht, Germany; 2KHz). The same A/D card was used to start the GVS and simultaneously record the EMG signals (except for during the walking experiment).

From the EMG signals from the erector spinae, the GVS artifacts (shifts due to voltage) were removed *a posteriori* by digitized data shift compensation. During the walking experiment, the presence of GVS artifacts on the EMG traces (voltage shift) was automatically detected (and then controlled for by subsequent visual inspection) on the left C7 erector spinae signal (the recorded muscle nearest the GVS electrode). During the other experiments, we simultaneously recorded EMG and a copy of the GVS stimulator output to precisely and unambiguously identify the GVS artifacts. For each EMG channel, the drift observed in the 40 ms following the GVS onset was modeled by a third order polynomial and removed from the signal. The same method was simultaneously applied to remove the artifact at the end of GVS. Then, the EMGs were numerically high-pass filtered (zero-phase Butterworth filter at 30 Hz), rectified, and smoothed with a 20 samples moving average.

For each stimulation, a 4 s time window was defined on both sides of the GVS onset (2 s before and 2 s after). For each subject, the EMGs were subsequently synchronized and averaged for each GVS polarity and condition and normalized by the activity measured before the occurrence of GVS (tonic activity during normal standing = 100%). The detection threshold to identifying VEMPs during the 300 ms following the GVS start was two standard deviations from the baseline. The baseline was defined as the median EMG during the 500 ms that preceded the GVS start. The medium latency of the VEMPs (ML; Fig. 4; Britton et al., 1993) was visually identified by comparing the EMGs following GVS to those at baseline or comparing the EMGs following GVS to those from opposed polarities if any.

The area of the VEMPs was computed by trapezoidal numerical integration of the rectified EMGs. The portions of EMGs used to compute the areas for each muscle in each subject in each experiment were the portions between the starting and ending time of the identified VEMPs averaged across all subjects. Then, we could compute the EMG areas in portions where VEMPs were expected, even if VEMPs were absent, allowing a between conditions comparison.

### Center of pressure

Static balance was assessed in a laboratory using a force plate (AMTI BP600900 with 6 degrees of freedom), and the position of the center of pressure was recorded at a 50 Hz sampling frequency. The data from the force plate were synchronized to the GVS start and averaged by subject, condition and GVS polarity.

### Kinematic recordings

We used a 3D-motion analysis system for the body segment analysis (Natural Point, Corvallis, USA). Twelve Optitrack Flex3 (640 × 480 pixels, 100 Hz) cameras were used in the gait experiment, and five Optitrack S250e (832 × 832 pixels, 250 Hz) cameras were used in the parabolic flight experiment. Prior to the data collection, calibration was performed to determine the camera orientation in relation to the working volume and the relative position of each camera to the other camera. Spherical reflecting passive markers (15 mm in diameter) were positioned on the head, trunk and body segments (see below). The data acquisition was performed using custom built software developed in MATLAB R2013a (MathWorks, Natick, MA, USA), enabling the simultaneous collection of synchronized kinematic and EMG data.

### Experiment 1: GVS during walking

Eleven subjects participated in the experiment (5 females and 6 males, 29.1 [±7] years old). The subjects walked on a treadmill (Pulsar 4.0, HP-Cosmos, Nußdorf Germany) at 0.75 m.s^−1^ with their eyes closed and gaze straight ahead. The subjects performed a 10-minute training before the first GVS stimulation. The subjects were secured by a harness, and the treadmill automatically stopped in case of falls in the harness (which never occurred). Each subject performed 12 trials, and each trial lasted 90 s with 5-minute pauses between each trial. During the experiment, each subject received more than 160 GVS (inter-subject mean=172) with unpredictable delay (>6s) and pseudo-randomized laterality (anode right or left).

We placed three reflective markers on each of the following segments: head, scapular girdle, pelvic girdle, hands, and feet. A real-time kinematic measurement of foot displacements was performed using a custom software developed with MATLAB to determine the phase of the gait cycle during which GVS occurred. Using this setup, we detected heel strikes based on the antero-posterior velocity of each foot. On the treadmill, the foot velocity shifted from positive to negative at the heel strike. GVS could be delivered at this time event with a latency as short as 40 ms (approximately 3% of the gait cycle). For each subject, the average gait cycle duration was determined during a ten-minute training, allowing GVS to be applied during the following four different phases of the cycle: (1) the first double support phase (immediately after the right heel strike); (2) the left swing (15% after the right heel strike); (3) the second double support phase (immediately after the left heel strike); and (4) the right swing (15% after the left heel strike). During the data analysis, each locomotor cycle was identified based on the right heel strike. During the post hoc analysis, we checked that GVS occurred at the proper phase.

To compare the effects of GVS during different phases of the gait cycle, we synchronized all kinematic signals to the GVS start, and the data were averaged to obtain one measure per subject, segment, cycle phase and GVS polarity. The anode right vs. left GVS induced medio-lateral deviations in opposite directions with the same delay. We compared the averaged medio-lateral velocities at both polarities to visually identify the latencies of the kinematic deviation. The maximum magnitude of the medio-lateral deviation was measured during the 500 ms following the GVS start.

The EMG signals were analyzed by performing a cycle categorization according to the phase of GVS occurrence (right heel strike, left swing, left heel strike, right swing, or no GVS). To analyze the phase effect on the VEMPs, the data were synchronized and averaged to obtain one sample synchronized to the GVS start per subject, muscle, and phase of the cycle. Finally, the EMGs of each muscle from each subject were normalized and expressed as a percent of the median activity during the No-GVS gait cycles. The occurrence and latencies of the VEMPs were visually identified by comparing the EMGs following GVS at opposite polarities.

### Experiment 2. GVS during upright standing with or without upper body contact

Nine subjects participated in this experiment (5 females and 4 males, 25.3 [±4] years old). We measured the effects of GVS on upright standing subjects under the following two conditions: (1) “freestanding” in which the subjects had to stay upright with their head turned to the right and their eyes closed and (2) “head contact” in which the subjects remained in the same position but with their head slightly in contact with a wall mounted on wheels allowing for precise proprioceptive feedback regarding body sway but could not be used as a firm support. Each subject was submitted to bipolar GVS with the anodal stimulation randomly set on the left or right mastoid process. Each condition included 20 trials, and each trial included 4 GVS with randomized polarity and an unpredictable delay (>5 s). The condition occurrence was randomized throughout the test session. Each subject experienced 160 GVS (2 conditions × 20 trials × 4 stimulations).

### Experiment 3. GVS in the postural muscle of free-floating subjects

This experiment occurred in the Zero-G airplane (A300, Novespace, Mérignac, France) chartered by the Centre National d’Etudes Spatiales (CNES, France) during two parabolic flight campaigns (VP-92 and VP-95, 2011-2012; 6 flights in total). During each flight, 30 parabolas were performed, and two subjects were tested (15 parabolas per subject). Each parabola provided a time lapse of 22 s of zero gravity (Fig. 5A), which was preceded and followed by 20 s time lapse of increased gravity (1.8 g; Fisk et al. 1993). Due to the influence of scopolamine on motor control (Bestaven et al., 2016), the subjects were not medicated.

Twelve subjects were tested in the upright standing position with their heads turned to the right, anode on the posterior mastoid process, and vision occluded by a mask. GVS occurred during normogravity (4 stimulations during each steady flight period between the parabolas), the first hyper-gravity period (n=4), and micro-gravity (n=4), resulting in a total of 180 GVS. All GVS were delivered at the same polarity to trigger VEMPs in the gastrocnemius and back muscles (anode posterior). Because the feet usually leave the floor in micro-gravity, the subjects did not have any contact with their environment and were secured. Three conditions were tested. Under the first condition, designated “Free-floating”, a scapular harness on the upper body secured the subjects (n=6) with distended tether (Fig. 5A). This harness did not apply any force on the subject, except for in the case of exceptional excursion (usually prevented by an operator situated in front of the subject during the experiment). The results of the first campaign (i.e., microgravity VEMPs remained present in the trunk only) prompted us to add other conditions to confirm that the observed effects were not due to (1) the presence of the harness on the upper body and/or (2) the lack of feet contact and external support. Thus, during the second campaign (VP95), the subjects (n=6) were tested under a second condition designated “Foot-strapped”. The subjects were submitted to different gravity conditions without a harness while being efficiently linked to the floor with their feet strapped to soft pads. The straps and pads we used (Concept X Kitesurf) allowed the ankle to be actuated in all directions with very good proprioceptive feedback. The same stimulation and recording protocols were performed during the first and the second campaign.

The third condition, i.e., “Harness”, was a control condition that was performed at our laboratory to consider the influence of gravitational detection and/or spine loading on the VEMPs of the lower limbs. In this experiment, the subjects were sustained in an upright position in a pelvian harness with their feet 10 cm above the floor. As in microgravity, the subjects stayed upright with an unloaded lower limb, but proprioceptive, visceral and otolithic perception of the gravitational field was present. The subjects also had their heads turned to the right, their vision was occluded, and they received 40 GVS with the anode on the posterior mastoid process and unpredictable delays. The same subjects (n=9) were tested under the standing upright (experiment 2) and Harness conditions.

#### Kinematics

We placed three reflective markers on each of the following segments: head, scapular girdle, and pelvic girdle. The 3D data were captured by 5 cameras (Optitrack s250e) at 250 Hz. The EMG data were synchronized and averaged to obtain one sample synchronized to the GVS start per subject, segment, Gravity and condition. Based on the averaged data, we manually identified the latencies of the GVS induced deviation based on the antero-posterior displacement velocity (sagittal plane). We computed the maximal posterior sway during the 1500 ms following the GVS start.

### Statistical analysis

For each experiment, a repeated-measures analysis of variance was performed using the MATLAB R2017a “ranova” function, and a post hoc analysis was performed using the comparison test of Scheffé. ANOVAs were computed to analyze the EMG area and latencies and kinematic magnitude and latencies. For each experiment, all conditions were considered within factor with repeated measures, except for the Free-floating vs. Foot-strapped conditions, which were considered between-factor because not all subjects participated in both campaigns (one condition per campaign). To statistically compare the proportion of VEMP occurrence, a Z-test was computed.

## RESULTS

### Experiment 1: Effects of bilateral GVS pulse during gait

This experiment was designed to investigate whether there was a phase dependency on vestibulo-spinal inputs during gait cycles and whether vestibulo-spinal inputs to axial or lower limb muscles were differentially controlled during walking. The subjects maintained their gaze straight ahead with their eyes closed, and GVS created a roll illusion contralateral to the anode, which elicited head displacement ipsilateral to the anode (Fig. 2A). Such lateral displacements were observed in the upper body segments (head, scapula and pelvis) and the stance foot (grounded during GVS but deviated at the following step). By contrast, the swing foot was deviated in the direction opposite to the upper body parts contralateral to the anode. The displacement magnitude in the frontal plane in the more rostral parts was higher (Fig. 2B; head: 6.9 cm (SD .9); scapular girdle: 6.1 cm (SD .8); pelvic girdle: 3.9 cm (SD .6)). The feet were deviated with a magnitude of 2.6 cm (SD .8) in the swing foot and 6.1 cm (SD 1.2) in the stance foot. The opposite directions of deviation between the upper parts of the body and swing foot along with the increased deviation observed in the more cranial parts elicits a body rotation in the frontal plane with an axis located at the hip level.

As expected, we found an effect of the anode side (i.e., polarity) and cycle phase, resulting in a significant interaction among Segment × Polarity × Phase (F(12,48)=23.6, p<0.001). The post hoc analysis revealed that the magnitude of the deviation of the head, scapular and pelvic girdles did not depend on the cycle phase at which GVS was delivered. By contrast, we found significant differences (Scheffe with p<0.05) in the feet displacements according to the cycle phase at which GVS was delivered. More specifically, in the right foot, the GVS effect during the first double stance phase did not differ from the left swing (both loaded phases), and the GVS effect during the second double stance phase did not differ from the right swing (both unloaded phases). However, each loaded phase (first double stance and left swing) significantly differed from each of the unloaded phases (second double stance and right swing). The same findings were observed in the left foot with no difference between the first double stance and left swing (loaded phases), no difference between the second double stance and right swing (unloaded phases), and significant differences between phases originating from each of these two groups (loaded vs. unloaded). This effect was observed in both GVS polarities (anode on the right or left side) but with opposite deviations.

The latency values of the kinematic deviations differed in specific body parts according to the phase (significant Segment × Phase interaction, F(12,48)=5.3, p<0.001), but the post hoc analysis revealed significant differences in the feet only. When GVS was delivered, a deviation occurred in the grounded foot during the following toe off. Consequently, in both feet, the deviation latencies during the loaded phases (swing of the contralateral foot, 533 ms (SD 127)) were significantly higher than those during the swing phases (214 ms (SD 47)). Similarly, the deviation latencies of the grounded foot were significantly higher than those of the head (203 ms (SD 135)) in the same phase but did not significantly differ from the latencies of the pelvic girdle (343 ms (SD 113)) and scapular girdle (333 ms (SD 105)). Significant differences were not observed among the head, scapular girdle, pelvic girdle and swinging foot deviation latencies.

The muscle activities observed during walking without GVS are consistent with previously observed activities on a treadmill (DeSeze et al., 2008; see Fig. 3 A). The erector spinae at T7 and L4 present a double burst rhythmic activity recorded in the right and left side with one burst after each heel strike. The gastrocnemius only presented one burst per cycle at the end of the stance phase (when the triceps surae is propulsive). To determine the GVS effects, we measured the EMG area and the latency value of the response immediately following GVS. As the full statistical plan was 7 muscle levels × 2 sides × 4 gait phases × 2 GVS polarities, we simplified the model by averaging the right and left muscles at each level with respect to the GVS polarity and loaded foot laterality. For instance, right ES L4 with right anode GVS and left foot loaded was averaged with left ES L4 with left anode GVS and right foot loaded to obtain ES L4 activity with “GVS ipsilateral” and “stance contralateral”. This resulted in the statistical plan 7 muscles × 2 GVS polarities (Ipsi/Contra) × 2 phases (Loaded/Unloaded side).

Interestingly, we did not observe significant effects of GVS polarity on the VEMP areas of *ES* at the C7 and T7 levels in the *rectus abdominis*, *pectoralis major* and *tibialis anterior*. An effect of the side of the loaded foot was observed in *rectus abdominis* (F(1,10)=5, p<0.05, Ipsitalic>Contra) and was near significant at ES C7 (F(1,10)=4.1, p=0.07, Contra>Ipsi). A significant effect of the GVS polarity was observed in EMG area ES L4 (Fig. 3B upper panel) with higher muscle activities following ipsilateral GVS than contralateral GVS (F(1,10)=15.1, p=0.003) and no loading phase effect. In *gastrocnemius*, a significant interaction between GVS polarity and the loading phase was observed (F(1,10)=34.5, p<0.001, Fig. 3B lower panel). The post hoc analysis revealed an effect of GVS polarity (Contra>Ipsi) in the gastrocnemius area only during ipsilateral foot loading.

Regarding the VEMP latencies, we compared the onset of the VEMPs elicited by contralateral GVS to ipsilateral GVS (VEMPs with the same latencies but opposite effects on the EMG magnitude). This analysis was performed at the ES T7, L4, Gastrocnemius and TA muscles, where the effects of the opposed GVS were easily identifiable. The two-way ANOVA (4 muscles × 2 phases) showed a main effect of muscle only (F(3,9)=67, p<0.001) on the VEMP latencies. The post hoc analysis revealed that the erector spinae at T7 and L4 did not have significantly different latencies (34 ms (SD 7) and 41 ms (SD 10), respectively), while these muscle latencies were significantly shorter than the lower muscles latencies at gastrocnemius and TA (140 ms (SD 18) and 173 ms (SD 30), respectively, not different). Altogether, these results indicate that the GVS effect is strongly phase dependent in gastrocnemius but not modulated by the gait cycle in the upper part of the body.

### Effects of GVS on upright standing subjects

It has been shown that in seated subjects or subjects standing upright while bearing a support, vestibular evoked responses were no longer elicited in the leg muscles (Fitzpatrick et al., 1994). In the present experiment, we investigated whether the same sensory gating of vestibulo-spinal inputs occurred in a similar way in the axial and leg muscles. Therefore, we compared the GVS-induced responses in the absence or presence of a contact with support. A bilateral GVS with the anode on the right (anode back) or left (anode front) mastoid process was delivered with the head turned to the right and occluded vision. The GVS with the anode back induced a backward sway (Fig. 4A) with an activation of the erector spinae and gastrocnemius muscles (Fig. 4BC). The maximum amplitude of the center of pressure during this antero-posterior sway was reduced by head contact (−2.9 mm (SD .9) in freestanding and −.2 mm (SD 1.48) with head contact; N= 9 subjects; Fig. 4A left vs. right panel). The two-way ANOVA revealed a significant interaction between the effects of polarity and condition on the antero-posterior magnitudes of the COP excursions (F(1,6)=18.3, p=0.005). The forward sway induced by GVS with the anode-front was also reduced (1.4 mm (SD 1.2) in freestanding and 1.2 mm (SD 1.41) with head contact), but this effect was not significant. The delay of the occurrence of the maximum sway was not significantly affected by either the polarity or head contact with a mean value of 379 ms (SD 60).

Several responses were elicited by GVS in the recorded muscles. As previously reported (Nashner et al. 1974; Britton et al. 1993; Fitzpatrick et al. 1994), GVS with the anode back first elicited a response with a short latency (single arrows, Fig. 4BC left panel) in the erector spinae and gastrocnemius muscles and a medium latency response (double arrow, Fig. 4BC). The short- and medium-latency EMG responses were in opposite directions at the lower limb level only but not in ES. We focused on the medium latency response, which was larger in amplitude, and its direction was correlated with the observed pattern of whole body sway. The comparisons of the VEMP occurrences indicates that the contact condition only influenced the gastrocnemius response. The GVS-elicited VEMPs from the erector spinae (L4 level; Fig. 4B) were detected in 78% of the subjects under the freestanding-condition, and this proportion was not significantly reduced to 56% under the head-contact condition (Z=1, p=0.16). The GVS-elicited VEMPs from the gastrocnemius were detected in 100% of the subjects under the freestanding-condition (Fig. 4B), but this proportion was significantly reduced to 22% of the subjects under the head-contact condition (Z=3.38, p<0.001).

In the erector spinae (L4), the VEMP area was affected by the GVS polarity. As previously described, GVS with the anode back (Fig. 4B) elicited a short latency inhibitory response and a medium latency response of a large amplitude. By contrast, when GVS was delivered with the anode front, the medium latency response was dramatically decreased. The two-way ANOVA revealed a significant main effect of polarity on the erector spinae elicited VEMP area (F(1,8)=8.5, p<0.05) and no effect of condition or interaction. Interestingly, the same analysis of the VEMPs elicited in the gastrocnemius revealed an interaction between the GVS polarity and contact condition (F(1,8)=7, p<0.05). The post hoc analysis showed a strong effect of the GVS polarity under the freestanding condition with a large response in the gastrocnemius to the anode back GVS and a marked inhibition with the anode-front GVS (Fig. 4C). This effect was drastically and significantly reduced under the head-contact condition with responses representing only 28% of the VEMP areas recorded while freestanding.

The VEMP latencies were measured at ES-L4 and the gastrocnemius under the Freestanding condition. The repeated-measures ANOVA revealed significantly shorter latencies in the erector spinae (49 ms (SD 26)) compared with those in the gastrocnemius (89 ms (SD 16); F(1,6)=11.7, p<0.05). The latencies at ES were not affected by the head contact (F(1,5)=0.26, p=0.63; 51 ms (SD 22) in free standing and 47 ms (SD 35) with head contact).

Altogether, these results revealed that incongruent proprioceptive cues had a major effect on the VEMPs in the gastrocnemius but not ES back muscles.

### Effects of GVS in modified gravity

To challenge the functioning of the vestibulo-spinal system, we performed experiments in which gravity was modified. Under microgravity, otolithic afferents are lacking because their activation depends on the effects of gravity; hence, the postural adjustments maintaining equilibrium become functionally useless.

Figure 5B present the experimental design of the free floating conditions (see Materials and Methods section). The subjects were submitted to GVS during parabolic flights (Fig. 5A) with periods of normal gravity, hyper-gravity (1.8 g) and micro-gravity (0 g). Due to the limited number of available trials under these conditions, GVS was delivered with the anode back only. GVS induced a backward displacement of the upper body in the antero-posterior direction under the normal gravity conditions (Fig. 5C, solid line, sagittal plane), which was increased under the hypergravity condition (dash-dotted line; F(2,8)=2, p=0.19 with n=4 subjects) and disappeared under the microgravity condition (dotted line). Under the normogravity and hypergravity conditions, all subjects expressed VEMPs in ES-L4 and gastrocnemius, whereas under the microgravity condition, 100% of the subjects expressed VEMPs in the left ES-L4, 60% of the subjects expressed VEMPs in the right ES-L4, and no subjects expressed VEMPs in the gastrocnemius (significant decrease, Z=5.2, p<0.001). We found a significant main effect of gravity on the ES-L4 muscle area (Fig. 6A1; F(2,8)=6, p<0.05 for left ES-L4 and F(2,8)=5.8, p<0.05 for right ES-L4). The post hoc analysis only revealed a significant difference between the hypergravity and microgravity conditions in the left ES-L4 area. We also found a significant main effect of gravity on the gastrocnemius area in the free-floating condition (Fig. 6A2; F(2,8)=23.8, p<0.05 for left gastrocnemius and F(2,8)=25, p<0.05 for right gastrocnemius). The post hoc analysis revealed that in both sides, the gastrocnemius EMG area was drastically decreased under the microgravity condition (quite null) and enhanced under the hypergravity condition (all p-values <0.05) compared with that under the normogravity condition.

Finally, a one-way repeated-measures ANOVA of the latencies at ES-L4 under the free-floating condition did not reveal any significant differences among the latencies at the three levels of gravity (micro-, normo-, and hyper-gravity; F(2,4)=3.1, p=0.15; Table 2).

Under the foot-strapped condition, we tested GVS only in microgravity and normogravity. Measuring the occurrence of VEMPs under the foot-strapped condition also revealed that the VEMP occurrence remained stable at the ES-L4 level (from 86% in normogravity to 75% in microgravity, Z=0.54, p=0.29 in ES-L4 right; from 100% to 86%, Z=0.53, p=0.29 in ES-L4 left), while the VEMPs were totally abolished in the gastrocnemius in the absence of gravity (from 100% to 0%, Z=7.3 and p<0.001 in both side). There was no significant difference in the ES-L4 VEMP area between the normogravity and microgravity conditions (Table 1; F(1,6)=2.9, p=0.14 in left ES-L4 and F(1,6)=0.84, p=0.39 in right ES-L4), whereas a main effect of gravity on the gastrocnemius VEMP was observed (left gastrocnemius F(1,6)=10.3, p<0.05; right gastrocnemius F(1,6)=6.7, p<0.05).

**Table 1.**
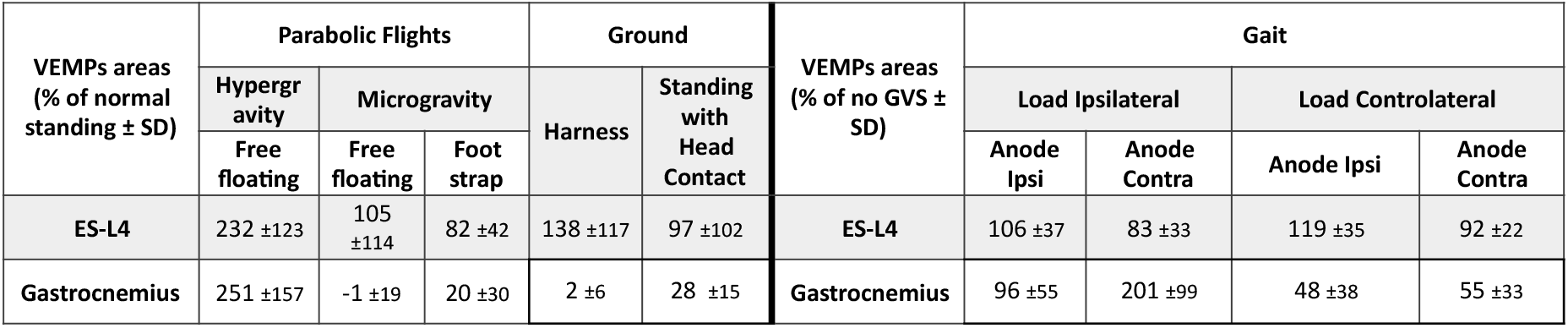
VEMP area under various conditions. Under the static conditions (modified gravity, harness, and head contact), the areas were normalized by the VEMPs observed under the adjoining control condition (i.e., 100% = VEMP area in free upright standing in normogravity). For the VEMPs during gait, the areas are expressed as a percent of the median activity measured during the gait cycles without GVS. Values represent the means ± SD.

| VEMPs areas<br>(% of normal<br>standing ± SD) | Parabolic Flights |  |  | Ground |  | VEMPs areas<br>(% of no GVS ±<br>SD) | Gait |  |  |  |
| --- | --- | --- | --- | --- | --- | --- | --- | --- | --- | --- |
|  | Hypergr<br>avity<br>Free<br>floating | Microgravity |  | Harness | Standing<br>with<br>Head<br>Contact |  | Load Ipsilateral |  | Load Controlateral |  |
|  |  | Free<br>floating | Foot<br>strap |  |  |  | Anode<br>Ipsi | Anode<br>Contra | Anode Ipsi | Anode<br>Contra |
| ES-L4 | 232 ±123 | 105<br>±114 | 82 ±42 | 138 ±117 | 97 ±102 | ES-L4 | 106 ±37 | 83 ±33 | 119 ±35 | 92 ±22 |
| Gastrocnemius | 251 ±157 | -1 ±19 | 20 ±30 | 2 ±6 | 28 ±15 | Gastrocnemius | 96 ±55 | 201 ±99 | 48 ±38 | 55 ±33 |

The repeated-measures ANOVA of the latencies with the condition (free-floating vs. foot-strapped) as a between subjects factor and muscle (ES-L4 vs. gastrocnemius) as the within subjects factor revealed a main effect of muscle (F(1,8)=37, p<0.001) with latencies at the ES-L4 (67 ms (SD 15)) that were much shorter than the latencies at the gastrocnemius (105 ms (SD 18)). Theses latencies were not significantly affected by the condition (see Table 2). Another two-way ANOVA was performed to analyze the latencies at ES-L4 only with the condition (free-floating vs. foot-strapped) as the between subjects factor and gravity (micro- vs. normo-) as the within subjects factor. Neither gravity nor the condition significantly influenced the latencies (F(1,6)=5.4, p=0.06).

**Table 2.**
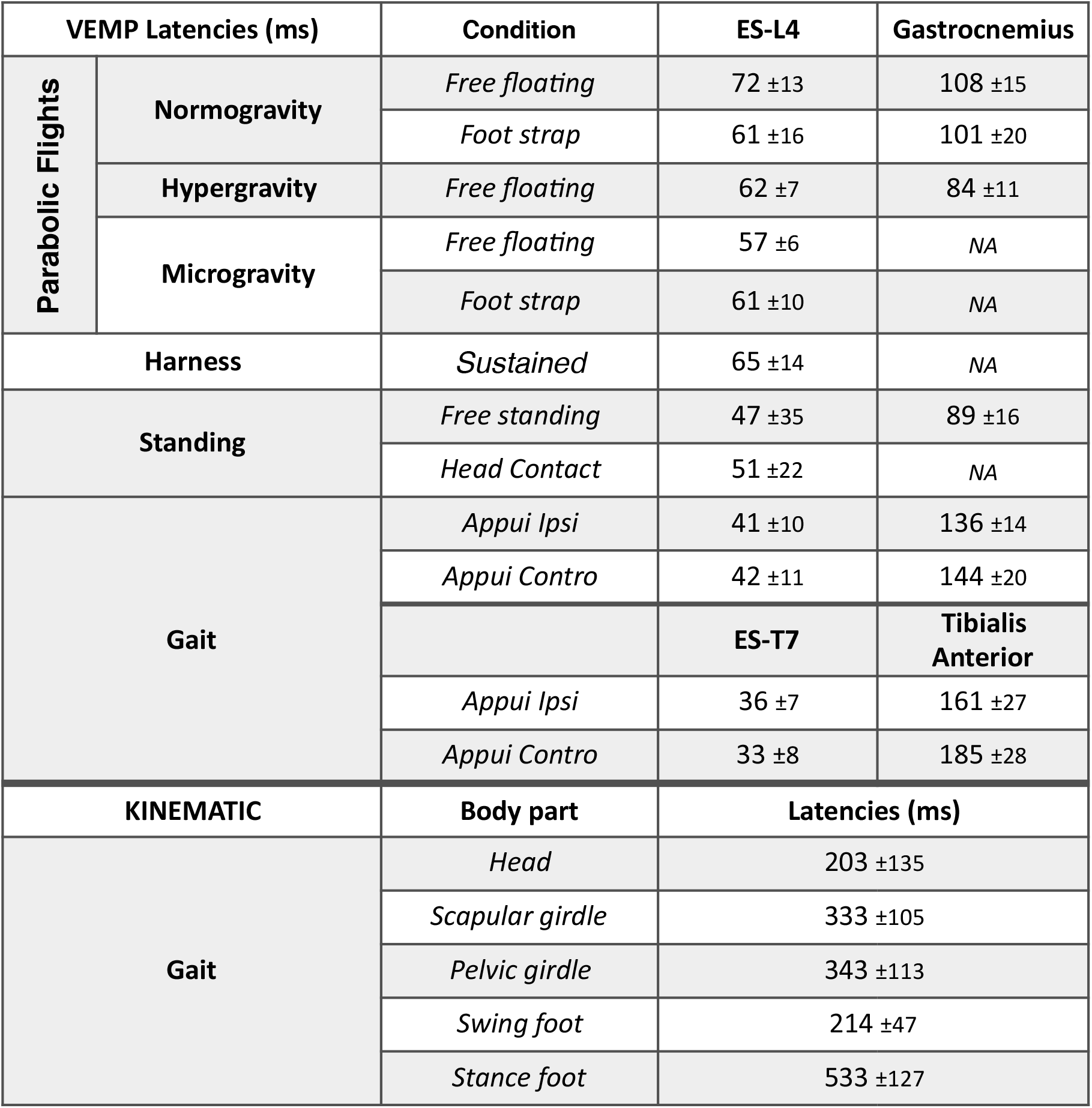
Averaged latencies of VEMPs measured in all experiments. Non-applicable (*“NA”*) is assigned when VEMPs were not observed under the corresponding condition. Values represent the means ± SD.

Finally, we performed a laboratory experiment in which the subjects were suspended in a harness. The aim was to evaluate the role of sensory cues in the gating of gastrocnemius VEMPs with subjects in the upright position, which is comparable to free-floating but with gravitational otolithic activation and spine load. The proportion of subjects in which GVS elicited VEMPs remained the same in the left ES-L4 prior to and during suspension (83%; Z=0, p=0.5), while there was a non-significant decrease (Z=0.67, p=0.25) in the right ES-L4 (from 83% to 67%). The VEMP area (Table 1) did not significantly decrease in the ES-L4 (F(1,5)=−0.4, p=0.7 right side, and F(1,5)=−0.9, p=0.4 left side). By contrast, the response to GVS completed disappeared in the gastrocnemius (from 100% of the subjects to 0% in the left gastrocnemius, Z= 3.5, p<0.001; and from 67% to 0% in the right gastrocnemius, Z=2.4, p<0.05) with a reduction in the VEMP area close to null in the gastrocnemius (F(1,5)=3.8, p<0.05 right side, and F(1,5)=2.7, p<0.05 left side). Altogether, the harness condition reproduced the same effects as those observed in micro-gravity with a stable proportion of subjects expressing axial VEMPs in the upright standing and harness conditions despite the gating of VEMPs in the gastrocnemius.

## DISCUSSION

We challenged the persistence of vestibular evoked myogenic potentials (VEMPs) in back muscles and leg muscle to determine how these muscles can be experimentally modulated. Our results strongly suggest that a differentiated control mechanism exists between axial and appendicular muscles when an imbalance is detected by vestibular inputs. The finding that with null gravitational information from the otolithic system (microgravity), we still observed persistent GVS elicited responses in axial muscles is particularly interesting.

### Functional responses to GVS in upright standing and phase-dependent effects during gait

The significant postural changes induced by GVS in upright standing (Fig. 1) and the mirror response observed when the anodal stimulation was delivered either to the right or left mastoid process (Fig. 3B & 4) validate the reliability of the stimulation protocol. This reliability was also confirmed under the various gravity conditions with a significant increase in the kinematic changes when the postural demand was increased (Fig. 5CII).

**Figure 1.**
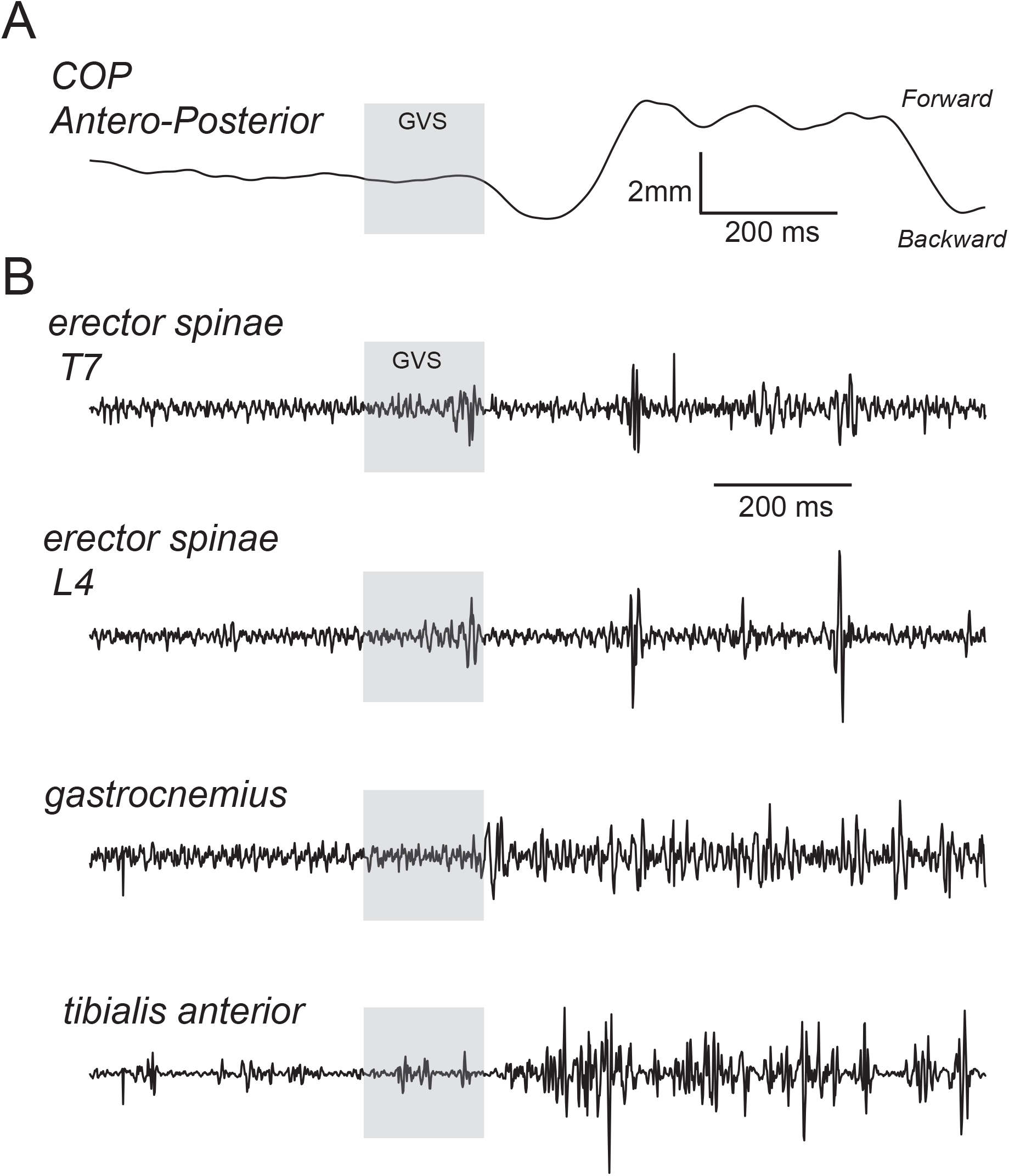
Representative kinetic and EMG recordings during GVS-induced perturbations. Recordings were performed in one subject with the head turned, vision occluded, and anode on the posterior mastoid. A, Backward displacement of the center of pressure. B, Electromyograms from (right) the erector spinae at the T7 and L4 levels, gastrocnemius and tibialis anterior.

**Figure 2.**
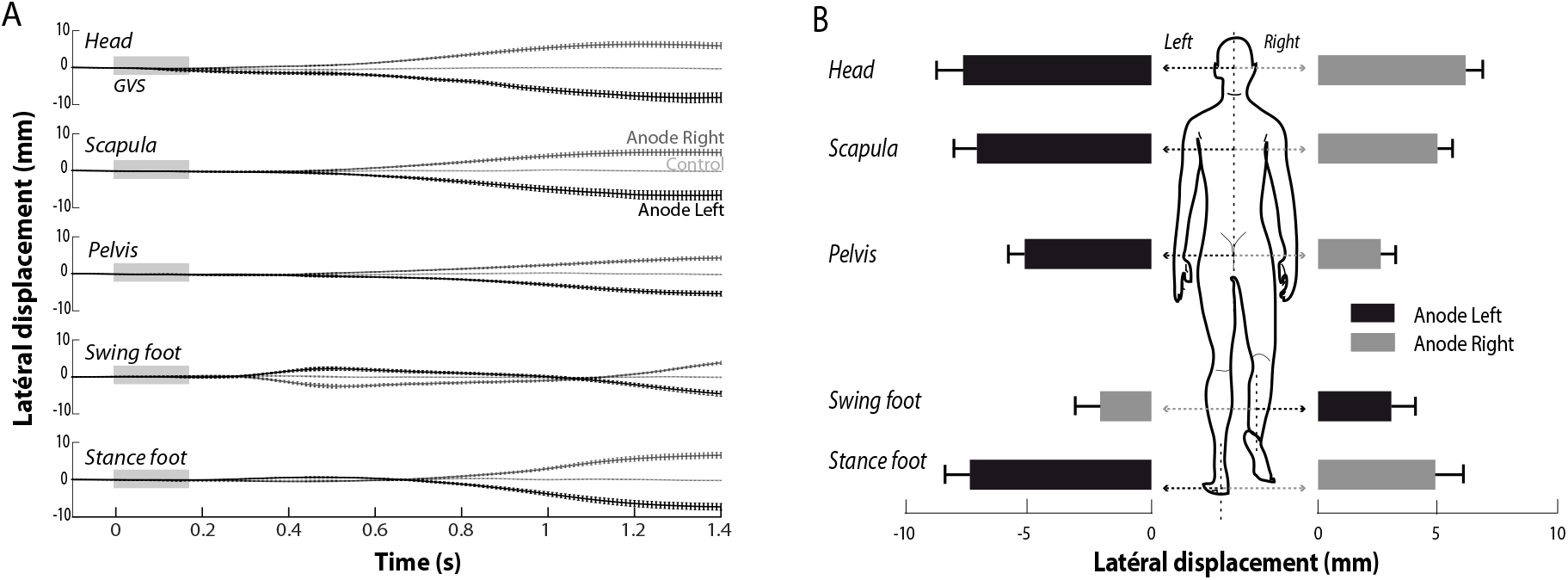
Rostro-caudal gradient of GVS-induced perturbation amplitudes during gait. A, Lateral displacements of the head, scapular girdles, pelvian girdles, and feet following GVS. Traces presenting data averaged across all subjects (N=11) with SD. The loaded stance foot following GVS was deviated during the following step, resulting in longer latencies than the swing foot. B, Magnitude of lateral displacements at various segmental levels. There was a decreasing gradient in the displacement magnitude from the head to the pelvic girdle. The swing foot was deviated in the direction opposite to the upper part of the body.

**Figure 3.**
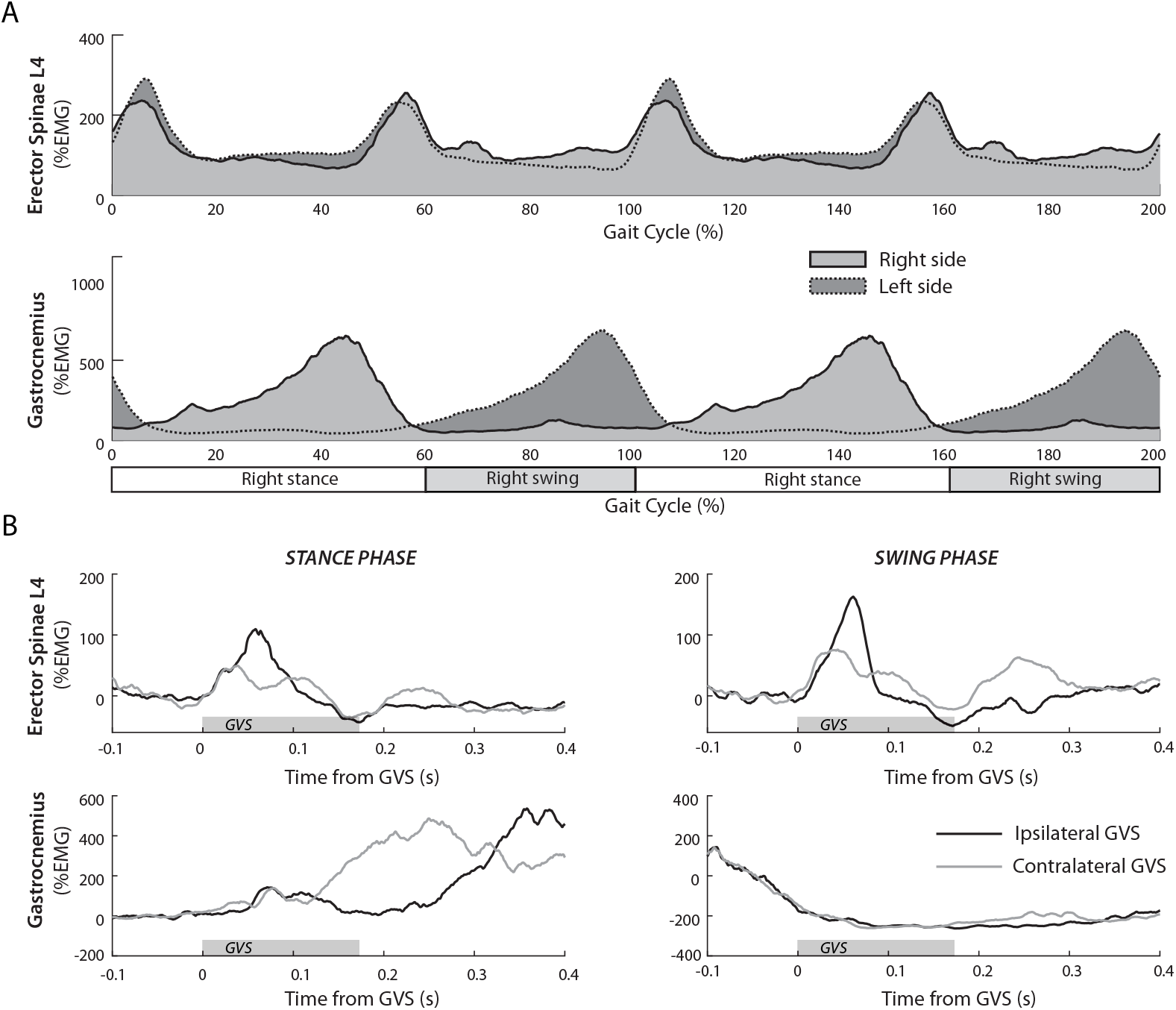
Phase-dependency of GVS-induced perturbations. A, Averaged rectified EMG of the right and left erector spinae (L4) and gastrocnemius during walking. Two successive normalized cycles (0-100%) are presented. ES muscles present bilateral double burst during each locomotor cycle, and each burst occurred close to the double support phase. Right and left gastrocnemius activity alternated with the burst occurring at the end of the stance period. B, The activity of the ES muscles was increased by GVS with the anode ipsilateral to the muscle and decreased with the anode contralateral to the muscle regardless of the phase of the cycle at which GVS was delivered. By contrast, the gastrocnemius activity was enhanced by the contralateral stimulation and decreased by the ipsilateral stimulation during the stance phase only and not during the swing phase, during which no GVS effect was observed.

**Figure 4.**
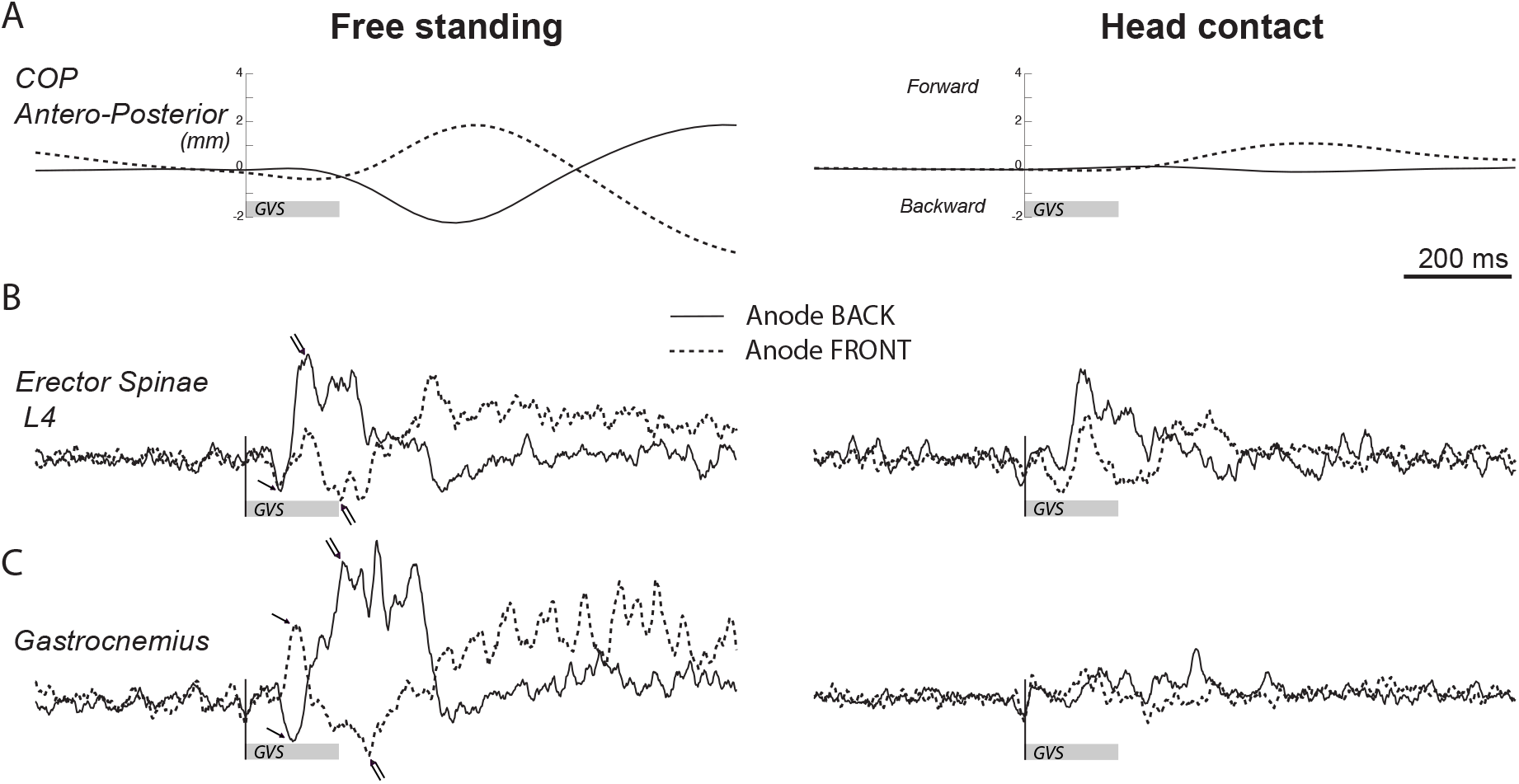
Maintenance of vestibulo-spinal reflex in axial muscles in upright standing subjects. A, During freestanding (left panel), the anteroposterior body sway was dependent on the GVS polarity (positive value = frontward COP displacement). The body sway was reduced close to null when the head slightly touched a support (tight panel, head contact). COP traces were averaged across all subjects (N=9). B, The medium latency response to GVS in the ES muscles was dependent on the GVS polarity and persisted in the absence (left panel) or presence (right panel) of a head contact. C, The medium latency response to GVS in the gastrocnemius muscle was dependent on the GVS polarity and virtually abolished in the presence (right panel) of a head contact.

**Figure 5.**
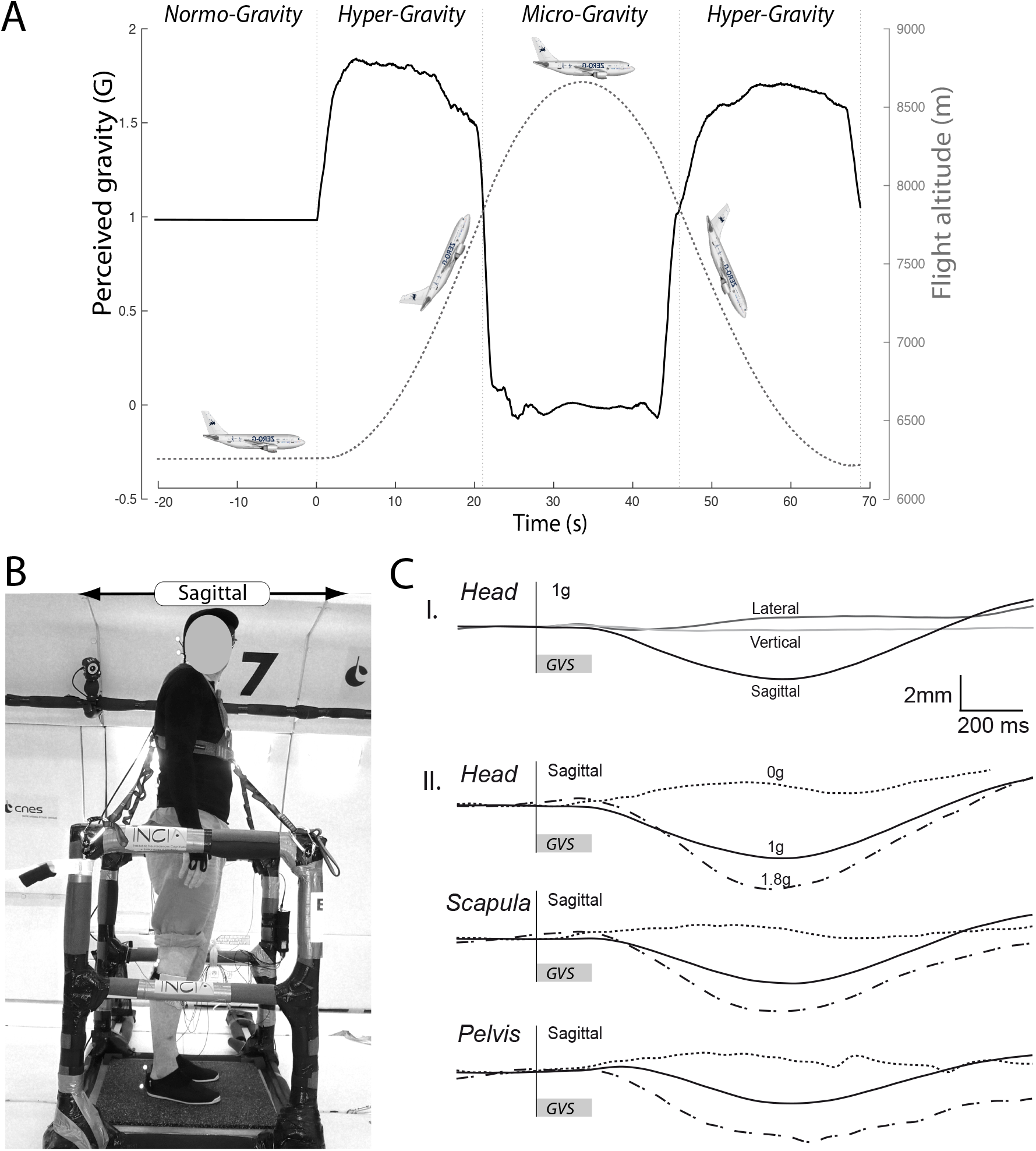
Experimental conditions during parabolic flight. A, Recordings from the plane altimeter (dashed line) and the accelerometer (black line, axis perpendicular to the plane ground) during one parabola. Each microgravity period (22 s) was flanked by two hypergravity periods (20 s), and the parabolas were separated by a steady flight period (>90 s). B, The subjects stayed upright with their vision occluded and head turned. The distended tether prevented the extreme excursion of the subject in microgravity. C, Displacements of the upper parts of the body following GVS with the anode on the posterior mastoid in (I) the lateral, vertical and sagittal plane of the head during steady flight and (II) the sagittal plane during steady flight (continuous line), microgravity (dotted line) and hyper gravity (dashed line) for the head, the scapular and the pelvian girdle.

**Figure 6.**
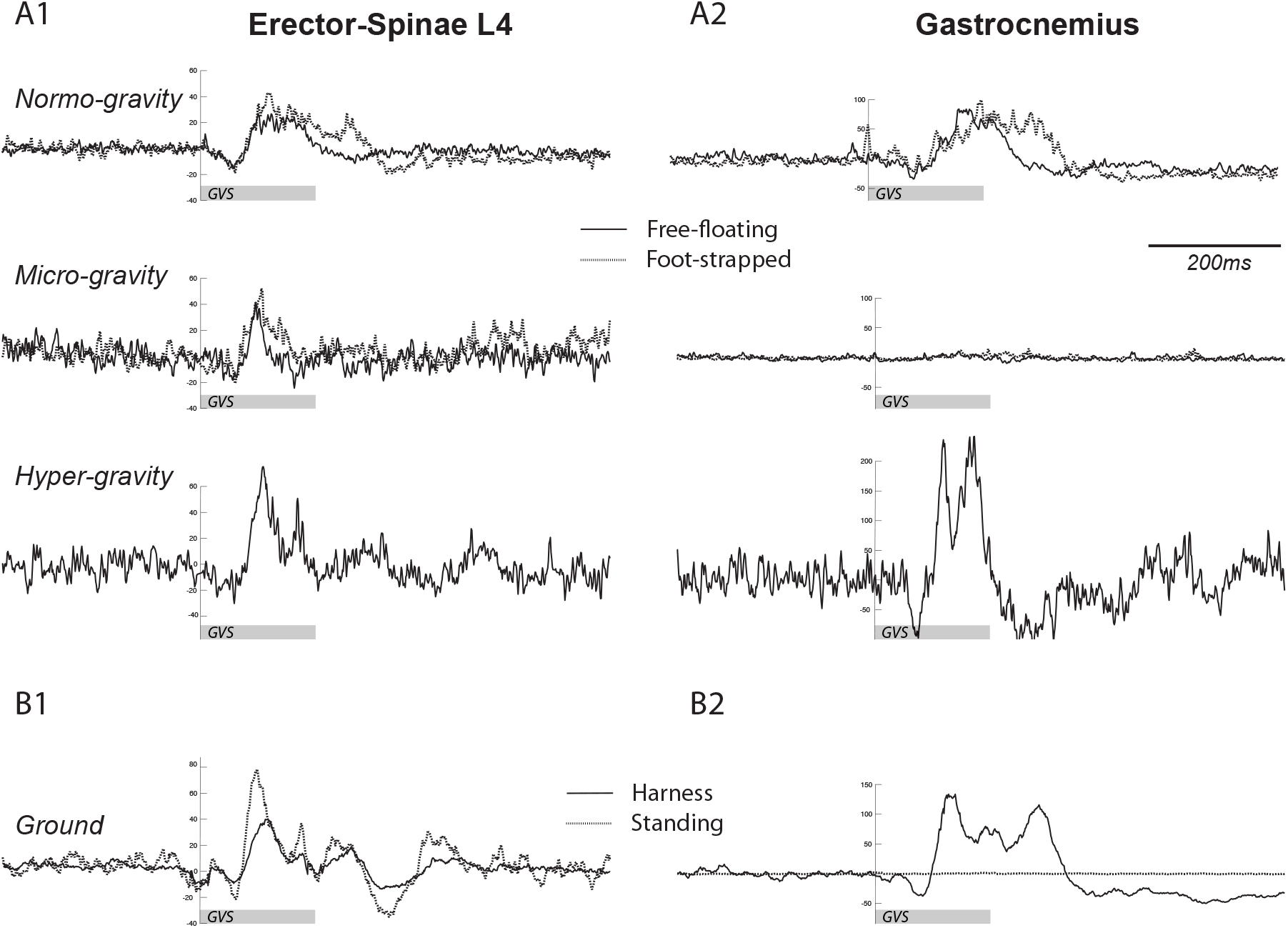
Maintenance of vestibulo-spinal reflexes in axial muscles with decreasing postural constraints. A, During upright standing, VEMPs (rectified EMG synchronized to the GVS start and averaged across the subjects) were elected by GVS (anode on the posterior mastoid process) from the erector-spinae L4 (left panels) and gastrocnemius (right panels) muscles. In microgravity, VEMPs were no longer observed in the gastrocnemius but were still present in the erector spinae. B, Under laboratory conditions, the VEMPs elicited by GVS from the erector-spinae L4 (left panels) and gastrocnemius (right panels) muscles were observed during freestanding in both the ES and gastrocnemius muscle but disappeared from the gastrocnemius muscles when the subjects were suspended upright in a harness.

During gait, the analysis of the lateral displacements following GVS indicates a global translation of the trunk to the anode side, i.e., the side opposite to the detected displacement, which is consistent with previous studies (Ali et al., 2003) and similar to the deviations observed with prolonged GVS (Fitzpatrick et al., 1999; Bent et al, 2000). The higher magnitude displacements in the upper parts of the body and the deviation of the swing foot in the reverse direction correspond to a body rotation in roll (Day et al., 1997; For a review, see Fitzpatrick and Day, 2004; Son et al., 2008) with a rotational axis located at the coxo-femoral articulation. During gait, VEMPs in the gastrocnemius were only recorded in the loaded leg, and the muscle discharge is increased when the loaded leg is ipsilateral to the anode, but the muscle activity is decreased with the contralateral anode. Similar phase effects were observed in the tibialis anterior (but with antagonistic action on the TA and GL; not presented). The phase-dependency reported here is consistent with the phase effects observed in gait kinematic (Bent et al., 2004; Dakin et al., 2013) and lower limb electromyography using stochastic GVS (Blouin et al., 2011) and the role of ground contact forces during unloaded gait (Ivanenko et al., 2002; Sylos-Labini et al., 2014). This modulation of gastrocnemius responses to GVS contrasts with the functional inflexibility of the ES-L4 muscle responses that remained constant along the gait cycle (Fig. 3).

### Differential gating of vestibular evoked myogenic potentials

One of our most striking observations is the clear dichotomy between the functioning of the vestibulo-spinal pathways impinging on either the axial or leg motoneurons. We observed a systematic suppression of VEMPs in the gastrocnemius when the force output did not contribute to the maintenance of body balance. This finding was observed (1) during the swing phase of the cycle during walking; (2) with head contact in the upright standing position; (3) in the absence of gravity to counteract; and (4) while the body was supported by a hip harness.

Experiments performed in modified gravity environments may help provide important insight into the sensorimotor integration of vestibular inputs and confirm that axial and leg muscles are differentially controlled. In microgravity, we reported a complete disappearance of VEMPs in the gastrocnemius muscle, in contrast to a major increase in the VEMP amplitudes in hypergravity to match the increasing destabilizing external forces. This VEMP deletion was not only observed during free-floating but also when cutaneous and proprioceptive feedback was provided through the ankle under the foot-strapped condition. Similarly, when the subjects were sustained in the upright position in a harness (Fig. 6B), i.e., a position similar to that adopted in the free-floating condition but with otolithic and proprioceptive knowledge of the context of normogravity, or stood upright with slight head contact (Fig. 4C), we still observed the deletion of VEMPs in the gastrocnemius. Therefore, the modulation of VEMPs in lower limbs is likely more related to body contact and support from the surroundings than to the postural attitude as observed in seated subjects (Britton et al., 1993; Day et al., 1997). The large decrease in the antero-posterior displacement of the center of pressure observed here (Fig. 4A), which has also been reported by other scholars (Britton et al., 1993; Fitzpatrick et al., 1994), has been interpreted as a gating of the vestibular input on the balance control, which could be dependent on the congruence of sensory feedback (visual, proprioceptive, and vestibular; Luu et al., 2012). Interestingly, when the subjects stood upright with slight head contact, a decrease was observed in the frontward and backward sway (both GVS polarities), demonstrating that this finding could not be explained only by the mechanical impossibility of sway against the forehead contact.

In contrast to the above discussion regarding the suppression of GVS-induced activity in the gastrocnemius muscle, a major result of the present study is the persistent responses in ES-L4 following GVS under all tested conditions. This persistence of ES-L4 VEMPs is particularly intriguing under the free-floating conditions in which the subjects were unplugged from their surroundings with no functional requirement of postural adjustments (no constraint of gravity on posture or balance and no visual frame of reference) without any body effectors available for support. The vestibulo-spinal responses observed in our experiments likely originated from the stimulation of the neuroepithelia of the cristae (in the ampula of the semi-circular canals), which was accompanied by concomitant stimulation of the maculae (Fitzpatrick et al., 2004). Thus, in microgravity, the otolithic signal was null, informing of the lack of gravity, and likely became incongruent when GVS occurred. In this context, the persistence of the axial response strongly suggests that the otolithic signal did not interfere with the integration of the semi-circular afferents. Together with the very short latency of the ES VEMP (47 ms versus 89 ms in the gastrocnemius in normal standing), the response of the axial muscles to vestibular inputs is more direct and robust and cannot be modulated by the context.

This dichotomy between the axial and leg responses to GVS could have functional significance considering the involvement of the trunk and ankle muscles in postural control. The interrelations between the body and its surrounding can be approached from the following two references values serving to control erect posture (Massion et al, 1998; Ivanenko and Gurfinkel, 2018) that can be dissociated in microgravity: one ‘geometrical’ value relative to the orientation of the body segment (“posture”) and one ‘kinetic’ value related to the distribution of the body mass with respect to the support area (“equilibrium”). Space flight experiments (Clément et al., 1984; 1988) have shown that the body schema, which is an internal representation of the body geometry, was surprisingly stable in microgravity, although extensive changes were observed in the labyrinthine and proprioceptive inputs. It can be hypothesized that axial VEMPs persisting in microgravity may originate from a process that supports the stability of the body schema. By contrast, Mouchnino et al. (1996) showed that the postural adjustment observed at the ankle during voluntary leg raising disappears in microgravity with an abolition of the shift of the center of mass (CM) toward the supporting limb. The variable VEMP we observed at the lower limb muscles could be related to the kinetic requirement of maintaining the center of pressure inside the polygon of sustentation, which is a constraint that disappears with the loss of gravity.

### Origin of the gating of VEMPs in the lower limb

Even if the exact mechanisms by which gating may occur remain unclear likely because multiple processes may be involved, it is a ubiquitous property of most vestibular responses. For example, the vestibulo-ocular reflex (VOR) can be suppressed during voluntary gaze behavior directed toward a visual target (Laurutis and Robinson, 1986; Pelisson and Prablanc, 1986). Similarly, the vestibulocolic reflex, which stabilizes the head in space by generating a command to move the head in the opposite direction (Colebatch et al., 1994), can be suppressed by voluntary head motion (Ezure and Sasaki, 1978; Goldberg and Cullen, 2011).

Regarding the suppression of VEMPs in the leg muscles, several mechanisms can be suggested to explain the gating. If during free floating, the persistence of VEMPs in the lower-limb muscles can be considered behaviorally irrelevant to maintaining balance, under the foot-strapped condition, the VEMPS also disappeared although there was direct active involvement of the ankle muscles. This finding suggests that gating is partially based on the detection of a gravitational field either by the otolithic system or extravestibular somatic receptors (Mittelstaedt, 1992). However, under the harness or head-contact conditions in normogravity, the detection of a downward gravitational field by upright subjects was not sufficient to elicit VEMPs in the gastrocnemius. Altogether, these results suggest that the incongruence among the vestibular, proprioceptive and cutaneous inputs abolishes the VEMPs in the lower limb muscles. The vestibular responses in the legs could be evoked only when all features of a nominal terrestrial postural task are met as follows: gravity must be detected, segment must be involved in the postural task, and sensorial cues must be congruent.

In their detailed study investigating the effects of vestibular stimulation during walking, Iles and colleagues (2007) reported that gating occurred in the soleus muscle during gait with similar phase-dependency as that observed in the present study, and these authors proposed that gating may occur through interactions with the central pattern generator at the lumbar level. Alternately, these authors also posed the hypothesis that gating could occur at the level of the brain stem. However, if this was the case, the axial and lumbar motoneuron responses could be affected in the same way since it has been recently shown in rodents that the two motoneuronal populations are similarly targeted by the lateral vestibulospinal tract (LVST; Kasumacic et al., 2010).

### Phylogenetic blueprint of vestibulo-spinal sensorimotor organization

The shift from swimming to upright bipedal locomotion is an outstanding feature of the evolutionary process, and the question of whether any of the underlying neural circuitries and mechanisms responsible for locomotor output production have been preserved during this transitional process remains unanswered. Several studies have previously suggested that during this evolutionary transition, the core features of the spinal cord circuitry of most primitive vertebrates, such as lamprey, have been conserved both in rats (Falgairolle et al., 2007; Beliez et al., 2015) and humans (de Sèze et al., 2008; Ceccato et al., 2009; for a review, see Falgairolle et al., 2006; Falgairolle et al., 2013). Nevertheless, the switch to terrestrial modes of locomotion and the emergence of limbs have necessarily increased the complexity of postural and locomotor control mechanisms, and currently, the trunk musculature is required for activity in strict coordination with limb movements during locomotion (Combes et al., 2004). Despite this more complex mode of coordination, the spinal networks responsible for trunk activity, even during bipedal walking, are still based on a bilaterally distributed spinal chain of oscillators as observed in lamprey (Grillner et al., 1995; Grillner and Wallen, 2002) but with the addition of limb CPG circuitry and associated interconnecting pathways. Using an amphibian model, several studies have explored the way in which during metamorphosis, locomotor networks and output transition from a purely trunk CPG (in the tadpole) to a limb CPG (Beyeler et al., 2008; for a review, see Sillar et al., 2008). Therefore, the extent to which the observations in the present study are related to comparable traces of primitive vestibulo-spinal pathways that could have been preserved in mammals is unclear.

Two main features of vestibulo-spinal responses may be highlighted in this context. First, the latencies of the vestibulo-spinal responses are on average twice longer in the gastrocnemius compared to those in the ES-L4 muscles (Table 2). Based on the typical conduction velocities reported in several studies (60 and 90 m/s for the lateral and medial vestibulo-spinal tract, respectively; Wilson et al., 1967; Akaike et al., 1973b; Akaike et al., 1973a), the theoretical delays required to recording the onset of muscle responses can be calculated. Assuming a 1 m length (from the head to the lower back) and a minimal conduction velocity of 60 m/s, including several synaptic delays, the vestibular inputs could elicit a response in approximately 30 ms in the ES-L4. In the gastrocnemius, this delay could be increased by up to 40 ms by adding the axonal route (approximately 1 m at 100 m/s) from the lumbar spinal cord to the muscle. Therefore, the dramatic differences between the latencies recorded in the ES-L4 and gastrocnemius cannot be explained by the sole differences in the target distances, suggesting that different pathways and/or mechanisms exist. However, it is unlikely that the differential gating operated on ES-L4 and gastrocnemius is attributable to the involvement of different pathways since the vestibular inputs to lumbar motoneurons are derived solely from the LVST in adult mammals (Wilson and Yoshida, 1969), and a recent studies (Graf, 2007; Kasumacic et al., 2010) demonstrated that only LVST establishes connections with both ipsilateral and contralateral thoracic motoneurons. As mentioned above, the possibility has been raised (Iles et al., 2007) that gating could occur at the spinal level. The emergent complexity of the circuitry organization required for hindlimb control could suggest that more complex local circuit interactions occur at the lumbar level. Recent studies have highlighted a major difference in the organization of mammalian neuronal circuits involved in postural and locomotor activities between the axial and lumbar level (Falgairolle et al., 2007; Beliez et al., 2015). Specifically, it has been found that neuronal activity propagates faster in the thoracic segments and that a strong slowdown occurs from the lumbar segments (Cazalets, 2005; Falgairolle and Cazalets, 2007). It has been suggested that as lumbar segments are intimately involved in hindlimb control, more complex local circuit interactions could slow the propagation through this region. The same vestibulo-spinal signal processing could occur at the lumbar level, which could explain the longer delay and serve as a basis for the sensory gating of vestibular inputs.

According to the phylogenetic perspective, the vestibular system has been present very early in most primitive vertebrates in a definite form sharing many common characteristics with the vestibular system in mammals (Kasumyan, 2004). This high degree of early development is likely related to the challenging situation faced by fishes in their three-dimensional permanently moving natural surroundings in which the sense of equilibrium plays an extremely important role in orientation, especially when vision is not involved. Detailed studies investigating the vestibular control of swimming in lamprey have characterized the commands sent to the spinal cord through reticulospinal neurons (Deliagina et al., 1992; Deliagina et al., 2006). This ancestral demand for highly efficient vestibular control could be the source of the strong hard-wired vestibulo-spinal inputs to axial motoneurons with a lower degree of flexibility than late emerging limb motoneurons.

## CONCLUSION

The results of the studies presented here are all suggestive of the differentiated governance of axial and appendicular segments when body rotation is detected by vestibular inputs. The persistent characteristics of the myogenic adjustments observed in the trunk even in absence of gravity suggest the presence of inflexible reflex wiring functionally efficient to maintain posture. However, the behaviors observed in the experiments during the transient microgravity episodes should be more related to the unconstrained expression of the neural circuitry adapted to terrestrial gravity than the response of adaptable processes to an unusual gravitational field. Multiple testing of postural behaviors in variable gravitational fields (lunar, Martian, etc.) could provide an opportunity to assess the level of the adaptability of these sensorimotor processes in new spatial contexts.

## Bibliographie

Ali AS, Rowen KA, and Iles JF. Vestibular actions on back and lower limb muscles during postural tasks in man. J Physiol 546: 615–624, 2003.

Beliez L, Barrière G, Bertrand SS, and Cazalets JR. Origin of thoracic spinal network activity during locomotor-like activity in the neonatal rat. In: Journal of NeuroscienceSociety for Neuroscience, 2015, p. 6117–6130.

Bent LR, Inglis JT, and McFadyen BJ. When is vestibular information important during walking? J Neurophysiol 92: 1269–1275, 2004.

Bent LR, McFadyen BJ, Merkley VF, Kennedy PM, and Inglis JT. Magnitude effects of galvanic vestibular stimulation on the trajectory of human gait. Neurosci Lett 279: 157–160, 2000.

Bestaven E, Kambrun C, Guehl D, Cazalets JR, and Guillaud E. The influence of scopolamine on motor control and attentional processes. PeerJ 4: e2008, 2016.

Blouin JS, Dakin CJ, van den Doel K, Chua R, McFadyen BJ, and Inglis JT. Extracting phase-dependent human vestibular reflexes during locomotion using both time and frequency correlation approaches. J Appl Physiol (1985) 111: 1484–1490, 2011.

Bresciani JP, Blouin J, Popov K, Bourdin C, Sarlegna F, Vercher JL, and Gauthier GM. Galvanic vestibular stimulation in humans produces online arm movement deviations when reaching towards memorized visual targets. Neurosci Lett 318: 34–38, 2002.

Britton TC, Day BL, Brown P, Rothwell JC, Thompson PD, and Marsden CD. Postural electromyographic responses in the arm and leg following galvanic vestibular stimulation in man. Exp Brain Res 94: 143–151, 1993.

Ceccato J-C, de Sèze M, Azevedo C, and cazalets j-r. Comparison of Trunk Activity during Gait Initiation and Walking in Humans. In: PloS one2009, p. e8193–8115.

Colebatch JG, Halmagyi GM, and Skuse NF. Myogenic potentials generated by a click-evoked vestibulocollic reflex. J Neurol Neurosurg Psychiatry 57: 190–197, 1994.

Combes D, Merrywest SD, Simmers J, and Sillar KT. Developmental segregation of spinal networks driving axial- and hindlimb-based locomotion in metamorphosing Xenopus laevis. J Physiol 559: 17–24, 2004.

Dakin CJ, Inglis JT, Chua R, and Blouin JS. Muscle-specific modulation of vestibular reflexes with increased locomotor velocity and cadence. J Neurophysiol 110: 86–94, 2013.

Day BL, Severac Cauquil A, Bartolomei L, Pastor MA, and Lyon IN. Human body-segment tilts induced by galvanic stimulation: a vestibularly driven balance protection mechanism. J Physiol 500 (Pt 3): 661–672, 1997.

de Sèze M, Falgairolle M, Viel S, Assaiante C, and cazalets j-r. Sequential activation of axial muscles during different forms of rhythmic behavior in man. In: Experimental brain research Experimentelle Hirnforschung Expérimentation cérébrale2008, p. 237–247.

Ezure K, and Sasaki S. Frequency-response analysis of vestibular-induced neck reflex in cat. I. Characteristics of neural transmission from horizontal semicircular canal to neck motoneurons. J Neurophysiol 41: 445–458, 1978.

Falgairolle M, Ceccato J-C, Seze Md, Herbin M, and cazalets j-r. Metachronal propagation of motor activity. In: Frontiers in bioscience2013, p. 820–837.

Falgairolle M, Falgairolle M, cazalets j-r, and Cazalets JR. Metachronal coupling between spinal neuronal networks during locomotor activity in newborn rat. In: The Journal of Physiology2007, p. 87–102.

Fisk J, Lackner JR, and DiZio P. Gravitoinertial force level influences arm movement control. J Neurophysiol 69: 504–511, 1993.

Fitzpatrick R, Burke D, and Gandevia SC. Task-dependent reflex responses and movement illusions evoked by galvanic vestibular stimulation in standing humans. J Physiol 478 (Pt 2): 363–372, 1994.

Fitzpatrick RC, and Day BL. Probing the human vestibular system with galvanic stimulation. J Appl Physiol (1985) 96: 2301–2316, 2004.

Fitzpatrick RC, Wardman DL, and Taylor JL. Effects of galvanic vestibular stimulation during human walking. J Physiol 517 (Pt 3): 931–939, 1999.

Forbes PA, Luu BL, Van der Loos HF, Croft EA, Inglis JT, and Blouin JS. Transformation of Vestibular Signals for the Control of Standing in Humans. J Neurosci 36: 11510–11520, 2016.

Forbes PA, Vlutters M, Dakin CJ, van der Kooij H, Blouin JS, and Schouten AC. Rapid limb-specific modulation of vestibular contributions to ankle muscle activity during locomotion. J Physiol 595: 2175–2195, 2017.

Goldberg JM, and Cullen KE. Vestibular control of the head: possible functions of the vestibulocollic reflex. Exp Brain Res 210: 331–345, 2011.

Goldberg JM, Smith CE, and Fernandez C. Relation between discharge regularity and responses to externally applied galvanic currents in vestibular nerve afferents of the squirrel monkey. J Neurophysiol 51: 1236–1256, 1984.

Graf WM. Vestibular System. In: Evolution of Nervous Systems, edited by Kaas JH2007, p. 341–359.

Iles JF, Baderin R, Tanner R, and Simon A. Human standing and walking: comparison of the effects of stimulation of the vestibular system. Exp Brain Res 178: 151–166, 2007.

Ivanenko YP, Grasso R, Macellari V, and Lacquaniti F. Control of foot trajectory in human locomotion: role of ground contact forces in simulated reduced gravity. J Neurophysiol 87: 3070–3089, 2002.

Kasumacic N, Glover JC, and Perreault MC. Segmental patterns of vestibular-mediated synaptic inputs to axial and limb motoneurons in the neonatal mouse assessed by optical recording. J Physiol 588: 4905–4925, 2010.

Kasumyan AO. The Vestibular System and Sense of Equilibrium in Fish. Journal of Ichthyology 44: S224–S268, 2004.

Laurutis VP, and Robinson DA. The vestibulo-ocular reflex during human saccadic eye movements. J Physiol 373: 209–233, 1986.

Luu BL, Inglis JT, Huryn TP, Van der Loos HF, Croft EA, and Blouin JS. Human standing is modified by an unconscious integration of congruent sensory and motor signals. J Physiol 590: 5783–5794, 2012.

Mars F, Archambault PS, and Feldman AG. Vestibular contribution to combined arm and trunk motion. Exp Brain Res 150: 515–519, 2003.

Mittelstaedt H. Somatic versus vestibular gravity reception in man. Ann N Y Acad Sci 656: 124–139, 1992.

Nashner LM, and Wolfson P. Influence of head position and proprioceptive cues on short latency postural reflexes evoked by galvanic stimulation of the human labyrinth. Brain Res 67: 255–268, 1974.

Pelisson D, and Prablanc C. Vestibulo-ocular reflex (VOR) induced by passive head rotation and goal-directed saccadic eye movements do not simply add in man. Brain Res 380: 397–400, 1986.

Smith CP, and Reynolds RF. Vestibular feedback maintains reaching accuracy during body movement. J Physiol 595: 1339–1349, 2017.

Son GM, Blouin JS, and Inglis JT. Short-duration galvanic vestibular stimulation evokes prolonged balance responses. J Appl Physiol (1985) 105: 1210–1217, 2008.

Sylos-Labini F, Lacquaniti F, and Ivanenko YP. Human locomotion under reduced gravity conditions: biomechanical and neurophysiological considerations. Biomed Res Int 2014: 547242, 2014.

Wilson VJ, Kato M, Peterson BW, and Wylie RM. A single-unit analysis of the organization of Deiters’ nucleus. J Neurophysiol 30: 603–619, 1967.

Wilson VJ, and Yoshida M. Comparison of effects of stimulation of Deiters’ nucleus and medial longitudinal fasciculus on neck, forelimb, and hindlimb motoneurons. J Neurophysiol 32: 743–758, 1969.

